# Sequential-GAM constructs the single-cell geometric 3D genome structure

**DOI:** 10.1101/2024.03.27.586762

**Authors:** Yongge Li, Kaili Wang, Minglei Shi, Michael Q. Zhang, Juntao Gao

## Abstract

Three-dimensional structure of the chromatin is crucial for cell identity and gene regulation. However, how genome-wide hierarchical geometric structure is organized in single cells remains elusive. Here we developed Sequential-GAM (Sequential Genome Architecture Mapping) to construct the hierarchical geometric structure and estimated the radial position of chromosomes, compartments, subcompartments, and genes in single cells by capturing contiguous thin sections of the nucleus. We found that several epigenomic features distributed gradually from the nuclear center to the periphery, including histone modifications and subcompartments. Besides, we estimated the variance of hierarchical structure and revealed the dynamic radial position of subcompartments B1 and B2. Furthermore, we defined the quasi-stable TADs set (q-stable TADs), in which TADs maintain a relatively stable distance to each other. Interestingly, q-stable TADs revealed the correlation between the stability of chromatin structure and chromatin activity. Finally, we discovered that the radial distance and the stability of radial positions for genes in single cells were negatively correlated with their expressions. Taken together, Sequential-GAM is able to estimate 3D genome’s geometric structure in single cell, and to reveal the structural stability and inter-cell heterogeneity.

## Introduction

In the nucleus, the genome is folded into hierarchical 3D structures. Previous techniques have been developed to investigate the topological 3D genome structure, including proximal-ligation dependent Hi-C^1^ and its derivatives, and proximal-ligation independent techniques such as genome architecture mapping (GAM)^2^ and split-pool recognition of interactions by tag extension (SPRITE)^3^. These studies have uncovered the hierarchical 3D organization of the genome, which includes chromosome territories, compartments, subcompartments, topologically associating domains (TADs), and chromatin loops^4,5^.

Due to the fact that the genome is a three-dimensional entity with a specific geometric shape, in addition to the topological structures mentioned above, genome function is also closely related to genome’s geometric characteristics, such as the area-to-volume ratio, sphericity, and radial position. For instance, during early development in mammals, it was found that the two X chromosome territories differed little in volume, but the active X chromosome territory showed a larger ratio of surface area to volume^6^. As for the radial position, when a gene transitions from inactive to an active state, it is likely that its radial position will change^7,8^. Active rRNAs are expressed within the nucleolus in the nuclear center, and heterochromatin is represented by nucleolus-associated domains (NADs) surrounding the nucleolus. Heterochromatin represented by Lamina-associated domains (LADs) is located around the nuclear periphery, and active transcription represented by mRNA is distributed between LADs and NADs^9^. The radial position of genes within the nucleus in a cell may also affect gene expression^10,11^. In addition to the above geometric properties, the radial distribution of chromatin is cell-type- and tissue-specific, changing during differentiation and development^12^. In addition, the shape of chromosomes changes from rod-like to more spherical between mitosis and late G1, which is related to chromosome decompaction^13^.

In order to comprehensively study the geometric features of genome, such as radial distribution, researchers developed GPSeq, which is an efficient method for examining radial structure with other chromatin markers for cell population. Nonetheless, GPSeq was incapable of resolving the radial chromatin arrangement and cell-to-cell viability in single cells^14^.

GAM^2^ is a proximal-ligation independent techniques to capture topological 3D genome structure. In principle, GAM measures chromatin contacts based on DNA sequencing from an extensive collection of random thin nuclear sections^2^. Thus, GAM technology enables the discovery of long-distance interactions and multiple interactions, driving substantial advancements in 3D genome research. Based on GAM, we hypothesize that by slicing and sequencing single cells layer by layer, we can obtain the exact information of the genome along a certain direction, then combined with constraints such as cLADs (constitutive LADs) and linear genome sequencing, we could construct the genome geometry of the single cell.

Therefore, based on the sequentially cutting of cell nuclear, we proposed Sequential-GAM to estimate the single-cell geometric structure and to investigate the radial position of genome-wide hierarchical structure. We utilized Sequential-GAM to investigate the radial distribution and structural variation of compartments, subcompartments, TADs, genes, and epigenetic features in the genome, which bridged the gap between geometry and topology of genome structure.

## Results

### Conception and simulation of Sequential-GAM

A basic idea of Sequential-GAM is that if we slice the nucleus sequentially, we could obtain the distances along a certain direction of different slices, which provides important information for the construction of chromosome geometry. The primary Sequential-GAM procedures are as follows. First, the cells or chromosomes in ground-truth were sectioned into 200nm thick slices, and the sequential parallel slices’ z-axis (perpendicular and pointing towards the top slice) position was determined thereafter. The radius and the relative position of the cuts were estimated, respectively, by the method of maximum likelihood (see Methods). Second, the loci were linked based on their linear position on the genome to calculate the genome distances. By the power-law relationship between chromatin 3D spatial distance and linear genomic distance^15^, we could estimate the distances among these loci. Third, three constraints were considered: (1) the z-axis information for each locus, (2) the estimated distance between each locus and cLAD (that is, the distance from the locus to the edge of the nucleus), (3) the estimated distance between every two loci. Finally, we calculated the geometric structure by minimizing the cost function under the constraints above, to estimate the coordinates of the locus in single cells (Fig. 1a).

**Figure 1.**
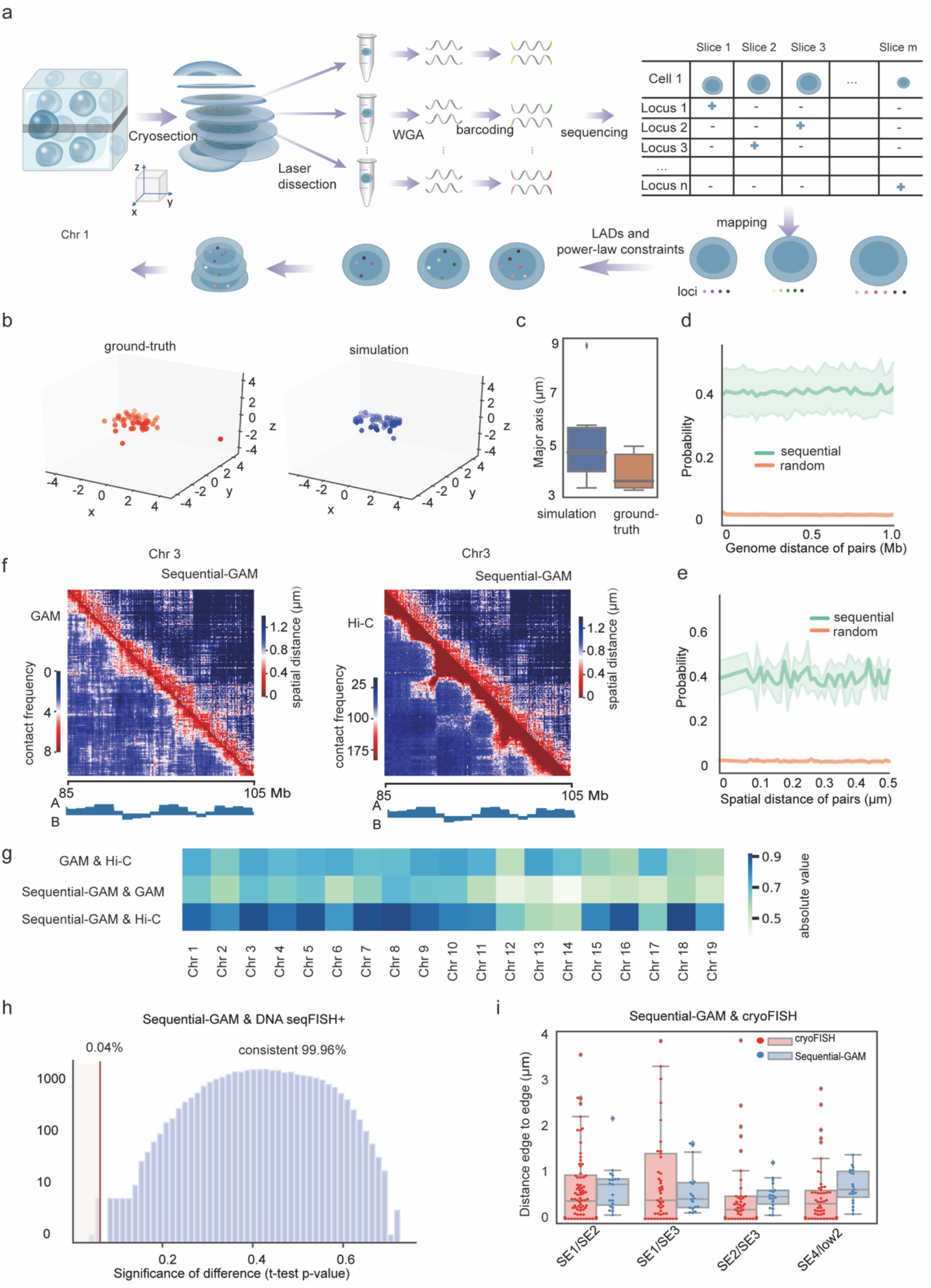
Sequential-GAM, a novel method to construct single cell 3D genome structure. **(a)** Schematic of Sequential-GAM workflow. Crosslinked cells (blue sphere) were embedded into gelatin and then dehydrated in sucrose. Next, cells were cryosectioned into the continuous 200nm-thick slices. Then, sequential thin slices were lined on the PET membrane slide. Nuclei were laser dissected on separated tubes, thereafter, and were further amplified and sequenced. The unique mapped reads in single cell are confined their positions by whether they are LADs and the power-law between the spatial distance of two loci and genomic distance. Then, the coordinated of z-axis of any loci is also used to constructed the three-dimensional structure of each chromosome. **(b)** The simulation of Sequential-GAM in Chr 1 from a cell based on DNA seqFISH+^16^ results. The red dots in the ground-truth (left) and blue dots in simulation (right) are arranged from the centromere (light color) to telomere (dark color). **(c)** The major axis of simulated and ground-truth in mES cells. **(d)** Comparison of the simulated probability of captured pairs with different genome distances using sequential slicing (red) and random slicing (green) method, separately. The simulations were performed in one structure of Chr 1 obtained from DNA seqFISH+^16^ data. The green lines represent the sequentially cutting and origin lines represent randomly cutting. The shadows represent the 95% confidence region. We obtained a 3D structure of one chromosome from DNA seqFISH+^16^, then randomly and sequentially cut the cell in different conditions, with 20 slices for Sequential-GAM in one cell and repeat 45 times with different cutting angles (to get 900 slices), and 900 times for GAM with random cutting angles. Then we compared the probability of 3600 paired loci captured in one cell simultaneously. **(e)** The difference of capture probability between sequential slicing and random slicing with various spatial distance. **(f)** Representative averaged distance matrix of distinct regions in Sequential-GAM and contact matrix in GAM and Hi-C at 100 kb bin. The Spearman correlation of Sequential-GAM over GAM and Hi-C is - 0.445 and -0.686 with p-values < 1e-5 at Chr3:85-105Mb, respectively. The color bar from red to blue indicates the observed spatial distance in Sequential-GAM (μm). The color bar in GAM means the contact probability. The color bar in Hi-C means the contact frequency. **(g)** The absolute value of Spearman correlation between Sequential-GAM and GAM or Hi-C for each chromosome at 1 Mb resolution. **(h)** The distribution of error of pair-wise distance for DNA seqFISH+^16^ dataset with Sequential-GAM. The p-values of the t-test of 29,903 pairs (99.96%) of the comparison were over 0.1; that is, the distances of pairs in Sequential-GAM were consistent with DNA seqFISH+^16^ dataset. **(i)** The comparison of the pair-wise distance for Sequential-GAM and cryoFISH. The p-values of the Welch’s t-test between DNA-FISH and Sequential-GAM were 0.108, 0.070, 0.179, 0.312, 0.069, separately, showing no significant difference.

To further estimate the reliability of the constraints in Sequential-GAM, we validated the Sequential-GAM algorithm in simulation data and in previously published chromosome structure^15,16^ (see Methods). This simulation was performed based on ground-truth reconstructed structure in 3D space, in which the theoretical power-law was applied (Supplementary Fig. 1a). Furthermore, the microscope-imaged single-cell structures of mES cells^16^ and human IMR90 cells^15^ were used for simulation (Fig. 1b, Supplementary Fig. 1b). Geometric characteristics, such as the structures and the chromosomal major axis, were consistent between ground-truth and simulation (Fig. 1b-c). Therefore, it showed that Sequential-GAM could reconstruct the genome structure with other independent published data.

Next, to investigate the advantages of Sequential-GAM over GAM, we simulated the random slicing in GAM and sequential slicing process in Sequential-GAM, by generating 900 slices from a genome structure in mES cells from DNA seqFISH+^16^. We found that the probability of captured loci in sequential sections was higher than that in random sections both in the linear genome distance and in the spatial distance, for different slice thickness and slice loss rates (Fig. 1d-e, Supplementary Fig. 1c-e).

### Experimental workflow and evaluation of Sequential-GAM

Sequential-GAM was applied in the mES F123 cell line, an F1 hybrid of *Mus musculus castaneus* × S129/SvJae with enormous public data. Briefly, cells were paraformaldehyde-fixed and embedded into gelatin at first. Then, cells were cryosectioned into continuous 200 nm-thick slices (Fig. 1a, Supplementary Fig. 2a). By tracking cell slices at specific locations, nuclei from the same cells were collected via laser dissection, and slices belonging to the same nucleus were collected into separate tubes (Supplementary Fig. 2b-d). Next, the DNA of each nuclear slice was amplified using single-cell whole genome amplification (WGA) and sequenced (Fig. 1a, Supplementary Fig. 2e-f, see Methods). We collected 128 nuclear slices from ten F123 cells in the G1 phase, ranging from 8 to 23 captured 200-nm slices in each cell (Supplementary Fig. 3a). The DNA within each thin slice was sequenced 5 million reads on average, and up to 61.6% coverage)of single-cell genome with a varied number of mapped reads was generated (Supplementary Fig. 3b, Supplementary Table S1).

Genome browser tracks of mapped reads from nuclear slices in the same cell showed that each nuclear piece contained a different fraction of chromosomes. For continuous nuclear portions in single cell, chromosomes showed preference on different slices in Sequential-GAM (Supplementary Fig. 3c-d). After filtering the noise (Methods, Supplementary Fig. 3e), we calculated the capture rate of the genome in single cells with the bin size of 1kb, 10kb, 100kb, and 1000kb, respectively, and chose 10kb as the bin size that was closest to the theoretical calculation (Methods, Supplementary Fig. 3f). Moreover, we found high correlations between the two batches of our experiments (Methods, Supplementary Fig. 3g).

Then, we separated the maternal (129S4) and paternal (CAST) genomes based on SNPs (see Methods, Supplementary Table S3). Next, we used cLADs sequences of F123 cells^17^ to constrain the chromatin fragments mapped to LADs. The position of fragments overlapped with cLADs were constrained to be within 0.5 μm of the edge within the nucleus. Then the positions of rest loci were calculated by the maximum likelihood estimation (see Methods) under the constraints of power-law relationship^15^ between chromatin 3D spatial distance and linear genomic distance. By this way, we calculated the geometric structure of each chromosome in single cells (Fig. 1a).

To evaluate the genome structure derived from the Sequential-GAM data, we compared it with the genome architecture produced by GAM^2^, Hi-C^18^ and DNA seqFISH+^16^. Considering that chromosome contact frequency can be viewed as a probabilistic description of the distance between two DNA regions in 3D space, we calculated the distance matrix using the Euclidean distance between each of the two DNA fragments in the three-dimensional structure (see Methods, Supplementary Fig. 4a, Supplementary Fig. 5a). Then, we compared the distance matrix from ensemble Sequential-GAM results and contact frequency matrix in GAM and bulk Hi-C with 100 kb bin. The measurements were highly consistent with those of GAM and of Hi-C (Fig. 1f-g, Supplementary Table S2). We also found a comparable relationship between genome distance and spatial distance, as well as comparable 3D structures in DNA seqFISH+^16^ (Supplementary Fig. 4b). To further validate the accuracy of the distance matrix, we compared the distance of 29,915 loci pairs in Sequential-GAM and DNA seqFISH+^16^ (Fig. 1h), 4 pairs in Sequential-GAM and cryoFISH^2^, respectively (Fig. 1i). The distance of 99.96% DNA pairs exhibited no significant difference between Sequential-GAM and DNA seqFISH+, all of which were comparable to cryoFISH. We also compared the 3D configuration in the allelic genome and found that Sequential-GAM revealed distinct organization of differentially expressed genes allelically. The differential spatial distances of gene TFPI2 in paternal and maternal showed differences associated with maternal allele expressed of TFPI2 in placenta-specific stage^19^(Supplementary Fig. 5b). Figure 1e shows that, according to simulation results, long-range hub could be obtained more efficiently by Sequential-GAM than by GAM (Supplementary Fig. 5c). In addition, we found that Sequential-GAM could identify the interactions between cis-regulatory elements (Supplementary Table S5-6). In summary, Sequential-GAM constructed the diploid single-cell genome structure of the mES cells.

### Radial organization of chromosomes and distinct genomic features

Next, we quantify the geometric features of each chromosome. It showed that the volumes of chromosomes increased along with the chromosome sizes (Pearson correlation is 0.785 with P-value as 6.881e-5 for paternal; Pearson correlation is 0.726 with P-value as 4.362e-4 for maternal. Fig. 2a, Supplementary Fig. 6a). In contrast, the ratio of surface area to volume increased in tandem with the decrement of chromosome size (Pearson correlation is -0.728 with P-value as 4.122e-4 for paternal; Pearson correlation is -0.468 with P-value as 0.0434 for maternal. Fig. 2b, Supplementary Fig. 6b). The sphericity of chromosomes is close to 0.514-0.923 in G1 phase (Fig. 2c, Supplementary Fig. 6c), similar to previous results in mouse ES cells^13^, as shown in the examples of chromosomes in Cell 8 (Fig. 2d, Supplementary Fig. 6d). In order to further understand the positioning of chromosomes, we also developed a method called Sequential-CT to calculate the relative position between two chromosome territories. By Sequential-CT, the relative positions of Chr 3, Chr 7, Chr 10 in Cell 9 were calculated and verified in DNA seqFISH+^16^ (Supplementary Fig. 6g, Supplementary Methods).

Next, we asked whether there is a radial preference of chromosomes territories. Here, we defined radial distance as the distance of a locus to the nuclear center. Although the position pattern of some chromosomes in the mES cell nucleus was revealed by chromosome painting^12,20^, the radial preference for all chromosomes in mES cells has not yet been explored. Thus, we calculated the average spatial distance for each chromosome to the nuclear center at 1 Mb resolution. Since different fractions were captured in different cells (Supplementary Fig. 6h, Methods), there would be experimentally induced bias if calculations were performed using the position of each locus of each chromosome in all cells. Therefore, we calculated the average radial distance for each chromosome in different cells. As shown in Figure 2e and Supplementary Figure 6e-f, the mean radial position of chromosomes positively correlated with chromosome size, while correlated inversely with average chromosomes’ gene expression, namely the chromosomes with smaller size and higher gene expression tend to locate closer to the nuclear center in mES cells. These results were consistent with prior studies in human embryonic stem cells^21^.

**Figure 2.**
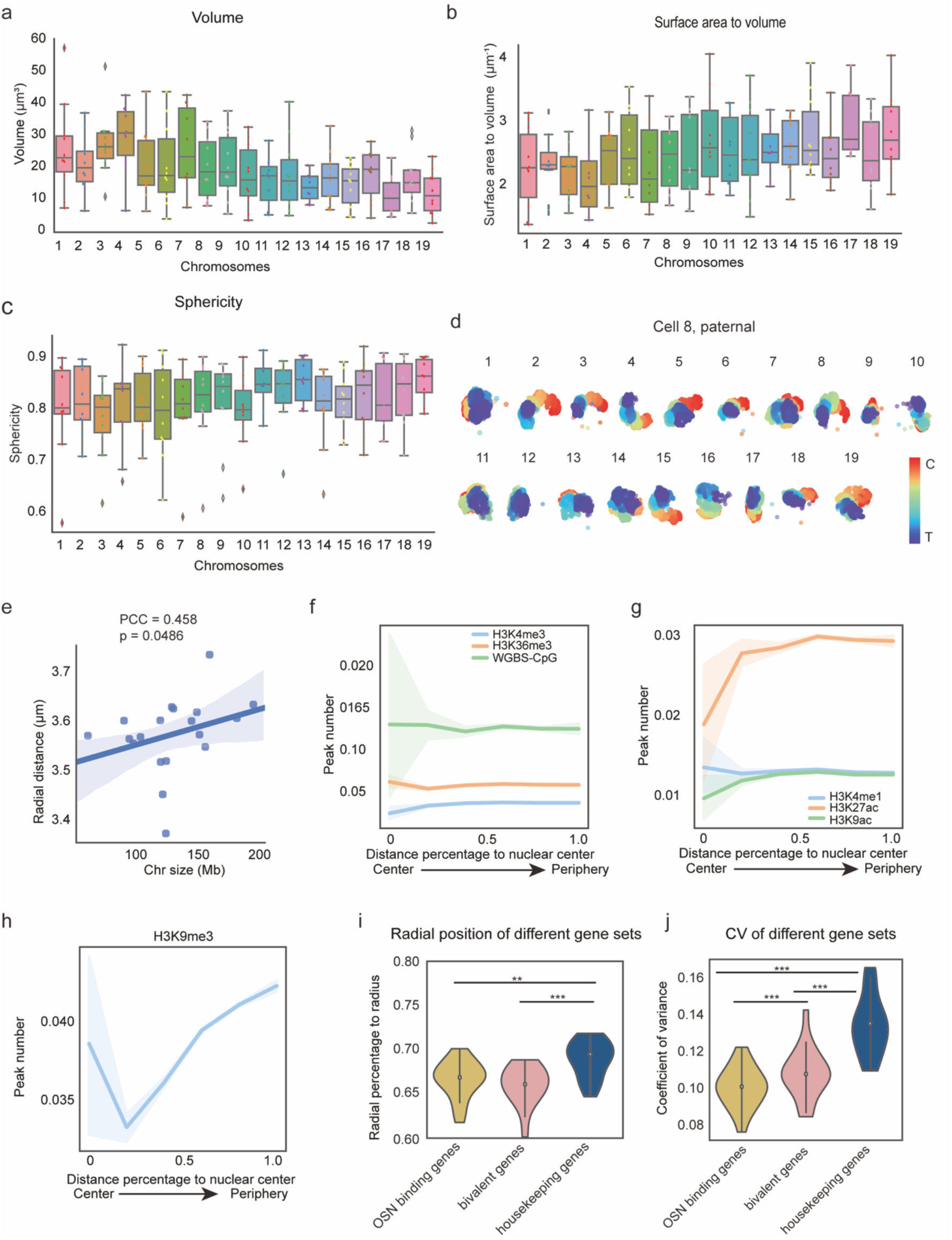
Radial organization of chromatin in mES cells. **(a)** The mean volume of individual paternal chromosomes. **(b)** The ratio of surface area to volume of each chromosome in paternal genome. (c) The sphericity of each chromosome in paternal genome in G1 phase. **(d)** 3D structures of paternal chromosomes in Cell 8 constructed by Sequential-GAM at 10 kb (each dot represent 10 kb), as an example. The linear genome is labeled as red (centromere) to blue (telomere). **(e)** The correlation between chromosome size (Mb) and radial distance to the nuclear center of each chromosome (um). Each dot represents the averaged value for a chromosome in 10 cells. The blue line showed the regression of the relationship between chromosome size and radial distance. **(f)** The radial distribution of epigenetic marks H3K4me3, H3K36me3, and DNA methylation CpG with 95% confidence at 10 kb. **(g)** The radial distribution of epigenetic marks H3K4me1, H3K27ac, and H3K9ac with 95% confidence at 10 kb. **(h)** The radial distribution of H3K9me3 with 95% confidence at 10 kb. (**i)** The radial distribution of OSN binding genes (green), bivalent genes(orange), and housekeeping genes (blue) in bulk cells. **(j)** The variance of the radial position of OSN binding genes, bivalent genes, and housekeeping genes in bulk cells marked by distinct colors. (The source data of Fig. 2f-j were in Supplementary Table S3.) (* indicate statistical significance as p<0.1, ** indicate statistical significance as p<0.05, *** indicate statistical significance as p<0.01, **** indicate statistical significance as p<0.001)

Genome structure is closely associated with gene expression, which is regulated mainly by epigenetic modification and transcription factors^22^. Therefore, we applied Sequential-GAM to measure the location of these features by integrating public data (Supplementary Table S3). As shown in Figure 2f, H3 lysine 4 trimethylation (H3K4me3) and H3 lysine 36 trimethylation (H3K36me3), which were enriched in promoter and gene body, respectively, exhibited little difference from the nuclear center to the nuclear periphery, while enhancer marks such as H3 lysine 27 acetylation (H3K27ac), H3 lysine 4 mono-methylation (H3K4me1) and H3 lysine 9 acetylation (H3K9ac) showed a mild trend of decrease (Fig. 2g), as well as the trend of RNA-seq (Supplementary Fig. 7a). The mildness maybe partially caused by the wide-spread position of these loci in the nucleus and the heterogeneous radial distribution among single cells^23^. On the contrary, the repressive marker H3 lysine 9 trimethylation (H3K9me3) showed a significant preference at the nuclear periphery than in the center (Fig. 2h). Similarly, the density of LADs and late replication regions increased gradually from the nuclear center to the periphery, while the early replication regions prefer to enrich at the center, compared with the late replication regions. (Supplementary Fig. 7a). In summary, the radial distribution of inactive epigenetic markers prefers continuous radial location along the nuclear radius, while active epigenetic markers show no significant preference.

Next, we asked whether different types of genes had preferred radial distribution. OCT4, SOX2, and NANOG (OSN) are master regulators of mES cells. Thus, OSN co-binding sites identified by ChIP-seq can be considered as cell-type-specific genes in mES cells^24^. OSN binding genes, bivalent genes, and housekeeping genes in different chromosomes appeared to have distinct radial preferences, along with the chromosomes (Supplementary Fig. 7b-c). Not like housekeeping genes, OSN binding genes exhibited similar positioning with bivalent genes (Fig. 2i, Supplementary Fig. 7b), indicating the similar radial position of fine-tuning gene regulation. To further investigate the radial position stability of different genes, we calculated the radial variance of each locus in ten cells. We found that housekeeping genes exhibited higher variance than OSN binding genes and bivalent genes, indicating the higher dynamics of housekeeping genes’ radial positions (Fig. 2j and Supplementary Fig. 7c). One reason for this might be the higher number of chromatin loops for cell-type-specific genes, which form spatially complex structure that constrains the structure change^25^.

Taken together, these results provide important insights into the radial preference and gradient distribution of chromosomes and distinct chromatin features.

### Dynamic positioning of chromatin compartments and subcompartments

Next, we examined the radial preference of compartments A and B in each cell. The cumulative distribution of compartments’ positions showed that compartments A located closer to the nucleus’ center, while compartments B located closer to the nuclear periphery (Fig. 3a-b). These results suggested the segregation of A and B compartments in a spatially polarized manner, indicating that the radial distribution of compartments is correlated with the function of chromatins, which is consistent with the previous studies^26^. We further divided the genome into subcompartment A1, A2, B1, B2, and B3, respectively, according to the correlation of their long-range interactions^27^ (more details in Supplementary Methods). The chromatin features of each subcompartment were well analogous with the previous study in human cells (Supplementary Fig. 8a)^27^. There is a gradient shift from the nuclear center to the periphery for subcompartments A1-B3 (Fig. 3c). Furthermore, we calculated the radial order of appearance for subcompartments from the nuclear center to the periphery, in which the order could represent the dynamic switch of radial position. We discovered that subcompartments A1, A2 and B3 were almost always located stably, that was, they always ranked 1^st^, 2^nd^ and 5^th^ in the order from the center to the edge of nucleus, respectively (Fig. 3d, Supplementary Fig. 8b). However, there were intersections between the radial order of B1 and B2, indicating the dynamic switch of B1 and B2 (Fig. 3d), as shown in the examples in Figure 3e-h. The phenomenon of such mix among different subcompartments was also observed in other reports^14^.

**Figure 3.**
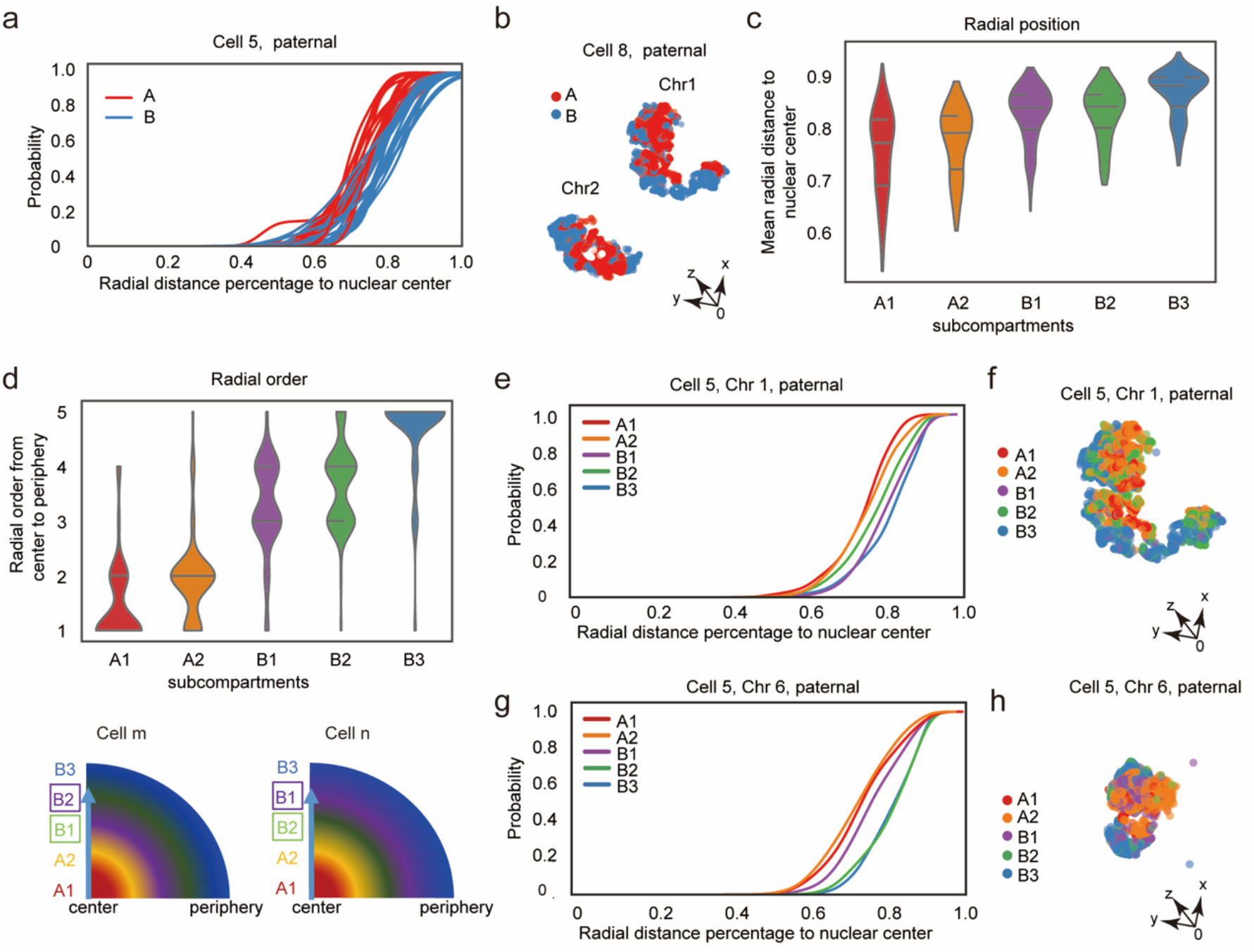
Radial organization of compartments and subcompartments. **(a)** The cumulative curves of radial distance percentage to nuclear center of compartment A (red) and B (blue) in all chromosomes in Cell 5, as an example. The radial distance percentage to nuclear center is obtained by calculating the radial distance divided by the radius of the cell nucleus. **(b)** The structure of compartment A (red) and B (blue) in paternal Chr 1 and Chr 2 in Cell 8 (each dot represents 10 kb. The coordinate axes are rotated compared to Figure2d). **(c)** The mean radial distance to nuclear center of subcompartments, which was normalized by nuclear diameter. **(d)** The radial order of subcompartments from nuclear center to periphery. 1 to 5 in y-axis indicate nuclear center to periphery. **(e. g)** The cumulative radial distribution of subcompartment A1 (red), A2 (orange), B1 (purple), B2 (green) and B3 (blue) in paternal Chr 1 on Cell 5 (e) and Chr 6 of Cell 5 (g). **(f. h)** The spatial structure of subcompartments in Chr 1 of Cell 5 (f) and Chr 6 of Cell 5 (h), with different subcompartments labeled by different colors.

Taken together, these results of Sequential-GAM provide independent evidence at the single-cell level, supporting the statement that there is a close correlation between the radial position of compartments and subcompartments and their transcriptional activity.

### The quasi-stable TADs set (q-stable TADs) reveals the relationship between genome stability and gene expression

TADs are regarded as the structure unit of genome with both spatial-temporal stability and evolutionary conservation^28,29,30^. However, when considering each TAD as a unit, a thorough understanding of the spatial stability of between TADs remains elusive. To further investigate the stability among TADs in spatial organization, we calculated the variance of the distance matrix for 6215 TADs in ten cells generated by Sequential-GAM (Fig. 4a). Next, we built the TAD graph by using TADs as vertices, the mean spatial distance among TADs as edges, and the variance as weight, of which we only preserved the edges with slight variance (Fig. 4a, Supplementary Fig. 9a-b). We then clustered TADs with relatively small variance insides by the Louvain algorithm^31^.

**Figure 4.**
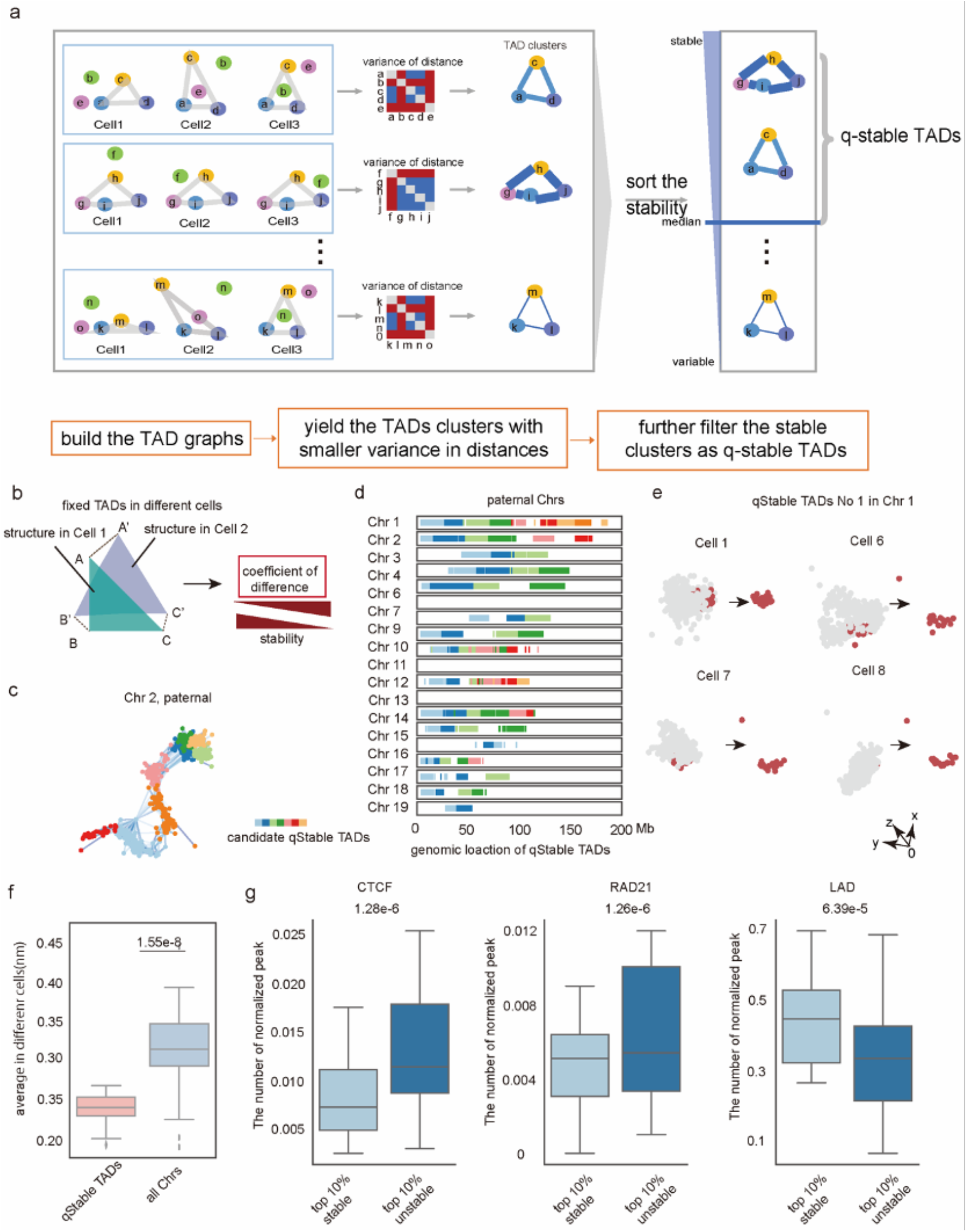
The q-stable TADs definition and stability in 3D genome. **(a)** The sketch map for the definition of q-stable TADs. The characters A-O represent different TADs in a chromosome. The colors in the variance matrix of TADs distance correlate with the distance of every two TADs. The darker blue means the smaller variance of two TADs among bulk cells. After getting the candidates of the q-stable TADs by Louvain algorithm from the distance matrixes, the q-stable TADs were sorted and filtered to get the stable part, with the coefficient of difference smaller than the median number. **(b)** Sketch map of coefficient of difference of two aligned structures representing q-stable TADs in different cells. The capital letters represent TADs. A, B, C and A’, B’, C’ represent TADs in different cells. **(c)** The graph of q-stable TADs candidates marked by different colored spots in paternal allele Chr 4. Each dot represents a TAD. The lengths of the lines between the spots represent the spatial distance between two TADs, and their colors represent the variance among ten cells; a greater variance results in a lighter color. **(d)** The linear position of the q-stable TADs in the paternal linear genome, which shows the locations of q-stable TADs with more than 3 TADs in the linear genome on different chromosomes. Different colors in each chromosome represents different q-stable TADs, while each stroke represents a 10kb loci in the genome. For different chromosome, the colors are repeated, and there is no relationship between the same color in different chromosomes. **(e)** 3D spatial structure of the q-stable TADs in different single-cells. In the left panel, the same q-stable TADs (red spheres) in Chr 4 (grey spheres) were shown in Cell 1, 8, and 10. In the right panel shows the No.3 q-stable TADs. Each dot represents a TAD. **(f)** The coefficient of difference of q-stable TADs and all chromosomes (p-value=1.55e-8). **(g)** The peak number of CTCF, RAD21, LAD and the gene expression level enrichment (Transcripts Per Million, TPM) in 99 stable q-stable TADs sets (coefficient of difference < mean -std) and unstable sets (coefficient of difference >= mean + std). The significances were calculated by Mann - Whitney U testThe p-value for CTCF, RAD21, and LAD are 1.27e-6, 1.26e-6, 6.39e-4, respectively. (* indicate statistical significance as p<0.1, ** indicate statistical significance as p<0.05, *** indicate statistical significance as p<0.01, **** indicate statistical significance as p<0.001)

To quantitively measure the stability of these TADs clusters, we defined a parameter called the coefficient of difference, which was the averaged distance between each TAD after the alignment of TAD clusters in ten cells (Fig. 4b, details in Supplementary Methods). Thereafter, we defined the part with the coefficient of difference smaller than the median as quasi-stable TADs set, termed q-stable TADs (Fig. 4a, Methods). The distances among TADs in the same q-stable TADs are relatively stable, while the distances among TADs in different q-stable TADs have higher variance. For example, eight q-stable TADs candidates were generated by Louvain algorithm in paternal chromosome 2 (Fig. 4c). After selected by coefficient of difference, six of these candidates were q-stable TADs, remained in chromosome 2 (Fig. 4d). Another example of one q-stable TADs visualized in different cells shows that the structures of q-stable TADs keep similar (Fig. 4e). In total, we identified 99 q-stable TADs in 3D space for the whole genome (Fig. 4d, Supplementary Fig. 9c-d). We also found that there were similar structures in DNA seqFISH+^16^ data, verifying the existence of q-stable TADs (Supplementary Fig. 9e). As computed in statistical analysis, the coefficient of difference of whole chromosomes was significantly higher than those of TADs in the q-stable TADs (Fig. 4f), suggesting that the q-stable TADs as structural units are more stable than the whole chromosome.

Next, we asked what correlated with the stability of these q-stable TADs. We obtained the most stable part and relatively unstable part from q-stable TADs (see Methods). The stable part showed lower percentage of CTCF and RAD21, which may be related to the structural dynamics due to loop extrusion, and lower gene expression level, but higher percentage of LAD (Fig. 4g). This suggested that topological structure proteins may play a role in q-stable TADs formation, which is coordinated with its function in TAD formation^32,33^. Overall, these results indicate that q-stable TADs are structural conformation that depicts the stability of the genome.

### The q-stable TADs reveal the association between structural variation and internal gene regulation

How geometric structural variability is related to gene regulation remains elusive. Therefore, we explored gene regulation in q-stable TADs. Firstly, we noticed a weak positive correlation between the coefficient of difference and averaged gene expression (Fig. 5a, PCC=0.22, p value= 0.027). Then we investigated the roles of specific genes or regulatory elements in q-stable TADs. We divided the q-stable TADs into three groups with four features (housekeeping genes, super enhancers, OSN genes and LADs), respectively (see Methods). The q-stable TADs enriched with super-enhancers demonstrate higher structural variation and higher gene expression than the q-stable TADs depleted of super-enhancers (Fig. 5b-c). These results indicated that the multiway interactions of super-enhancers in 3D nucleus space contributed to the instability of inter-TADs, as well as transcription ^2,34^.

**Figure 5.**
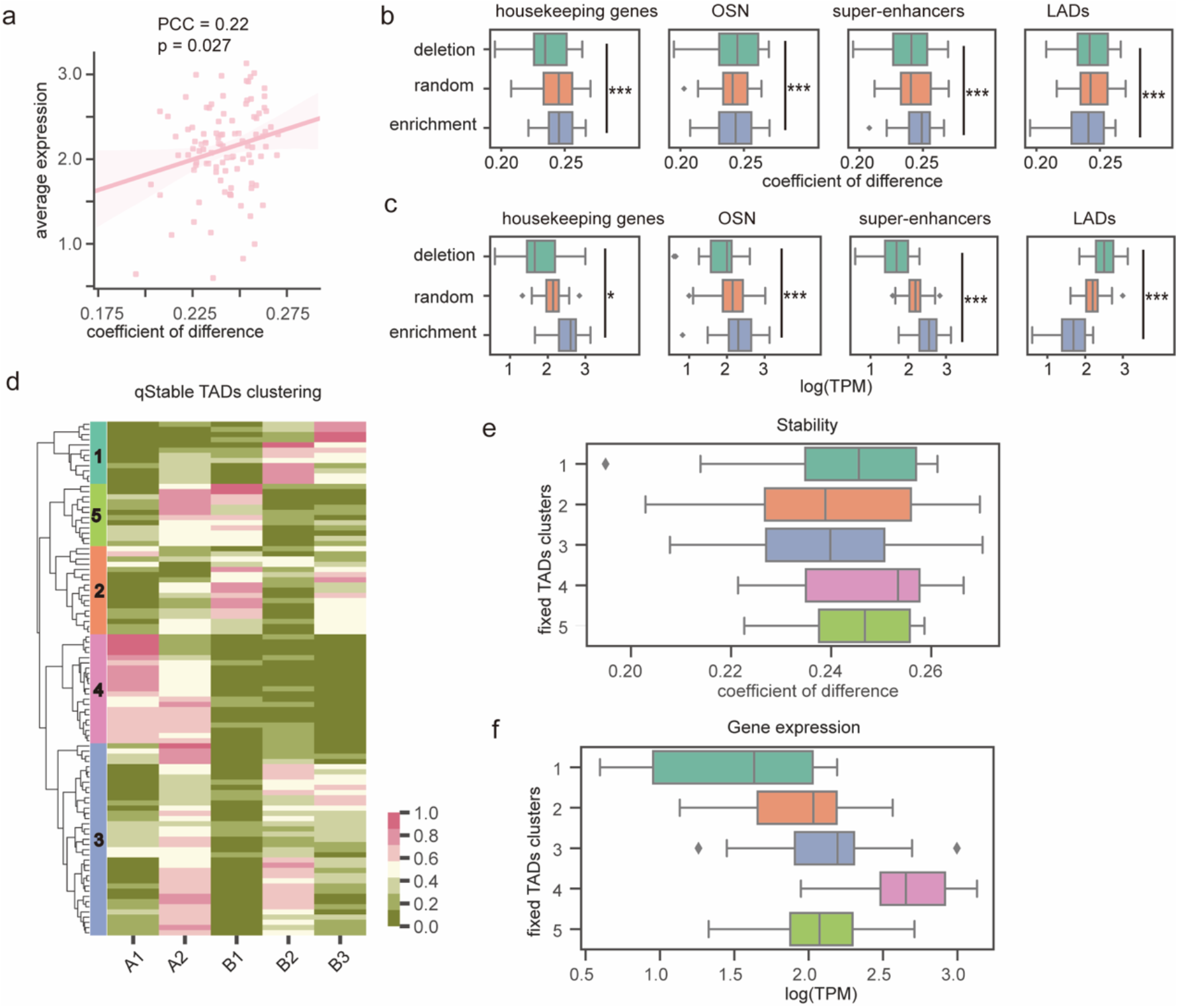
The q-stable TADs regulate gene expression. **(a)** The relationship between the mean expression of transcripts in the q-stable TADs and the structural variation of the q-stable TADs (represented by coefficient of difference), with the linear correlation shown in the pink line and the 95% confidence coefficient shown in the pink shadow. The number of q-stable TADs here are 191. **(b)** The coefficient of difference of all q-stable TADs which were enriched (mean peak num >= 0.7 quantile) or depleted (mean peak num < 0.3 quantile) with housekeeping genes, OSN-binding genes, super-enhancers and LADs, from left to right. The number of q-stable TADs in depletion, random, enrichment group for housekeeping genes is 41, 29, 29, respectively; for OSN-binding genes is 38, 36, 25, respectively; for super-enhancers is 38, 32, 30, respectively; for LADs is 36, 32, 31, respectively. **(c)** The gene expression in each group coordinated with (b). **(d)** The heatmap of q-stable TADs which enriched or depleted with subcompartments A1, A2, B1, B2, or B3 were clustered into six groups. The number of q-stable TADs in group from 1 to 5 is 12, 17, 37, 21, 12, respectively. **(e)** The coefficient of difference for each cluster in (d). **(f)** The gene expression for each cluster in (d) is labeled with different colors. The significances were calculated by Mann - Whitney U test (p values of figure 5e-f are in Supplementary table S4). (* indicate statistical significance as p<0.1, ** indicate statistical significance as p<0.05, *** indicate statistical significance as p<0.01, **** indicate statistical significance as p<0.001).

However, as mES-specific genes, the inverse pattern was observed in q-stable TADs enriched with OSN-binding genes and LADs (Fig. 5b-c), indicating the possible different roles of OSN binding genes and super-enhancers in 3D structural variation. The possible reason for the differences may be that the regulation of OSN-binding genes is typically important, so that they tend to have more complex regulatory regions which lead to a rather stable 3D structure^25^. The structural stability of the LADs-enriched q-stable TADs is higher, whereas the expression of theirs is lower (Fig. 5b-c), suggesting that LADs may serve as anchor point for the structure of q-stable TADs, and stabilize inter-TADs structures, consistent with LAD’s function in maintaining the mechanical structure of the nucleus^35^. These works indicated that the stability of q-stable TADs is related to internal chromatin features.

To further characterize the characteristics of all q-stable TADs, we performed hierarchical clustering with the enrichment of housekeeping genes, OSN binding genes, super-enhancers, and LADs, respectively. The cluster enriched with OSN-binding genes and LADs are significantly more stable than the cluster enriched with housekeeping genes and super-enhancers (Supplementary Fig. 10a-c), indicating that LAD, housekeeping genes and super-enhancers have distinct functions in the stability of q-stable TADs.

Since subcompartments reveal the chromatin state profoundly^27^, we investigated how structural stability was related to the subcompartments. In our study, we clustered all q-stable TADs based on the different subcompartment enrichment ratios into five groups, meticulously evaluated the structural stability of various chromatin compartment combinations, with a cursory examination of their influence on gene of compartment B in its combinations, especially when juxtaposed against interactions involving A compartments. In comparisons within A compartment pairings, A2+B2 exhibits a notably higher stability compared to A2+B1, indicating a preferred stable structure in the A2 and B2 association (Fig. 5e-f, Supplementary table S4). In contrast, A1+A2 shows less stability compared to A2+B1(Fig. 5e-f). This finding suggests a distinct stability trait inherent to the A2+B1 combination, despite both involving the A2 compartment. In summary, these results accentuate the complexity and significance of chromatin structural stability and its subtle yet potential impact on gene expression patterns.

### The radial distribution of genes correlates with their local status

At genome-wide scale, the radial variation and the activity of genes have not been explored in detail^14^. As gene positioning in the nucleus is determined not only by the gene itself but also by the neighboring genes or its interacting partners^36,37^, we calculated the spatial positions of genes with the bin of 1 Mb. The radial distance from nuclear center of genes was negatively correlated with their local transcription (Supplementary Fig. 11a). To further characterize the correlation, genes were categorized as active or inactive based on the compartment to which they belonged, and the radial position of each gene was computed. The radial distance to nuclear center of gene loci in compartment A was negatively correlated with the gene expression of that gene in each chromosome, while the radial distance of genes in compartment B showed a slightly positive correlation with gene expression (Supplementary Fig. 11b-f). This suggested that, at the genome-wide scale, the radial distance from nuclear center of genes was correlated with their transcription activity. Active genes located closer to the center of the nucleus, while inactive genes located close to nucleus periphery.

We also investigated the corresponding variation of genes’ radial location in 3D space among single cells (see Methods). Interestingly, the radial variation of gene loci in 3D nucleus was positively correlated with the local transcription levels (Fig. 6a). The genes in compartments A display greater radial variation, whereas inactive genes in compartments B exhibit smaller variation (Fig. 6b, Supplementary Fig. 12a-b). The radial position of genes in subcompartment A1-B3 exhibited a stepwise reduced correlation with their variation (Supplementary Fig. 12c-d). In summary, the radial locations of genes which are more active, are more structure dynamic among single cells, whereas the radial locations of inactive genes were more stable.

**Figure 6.**
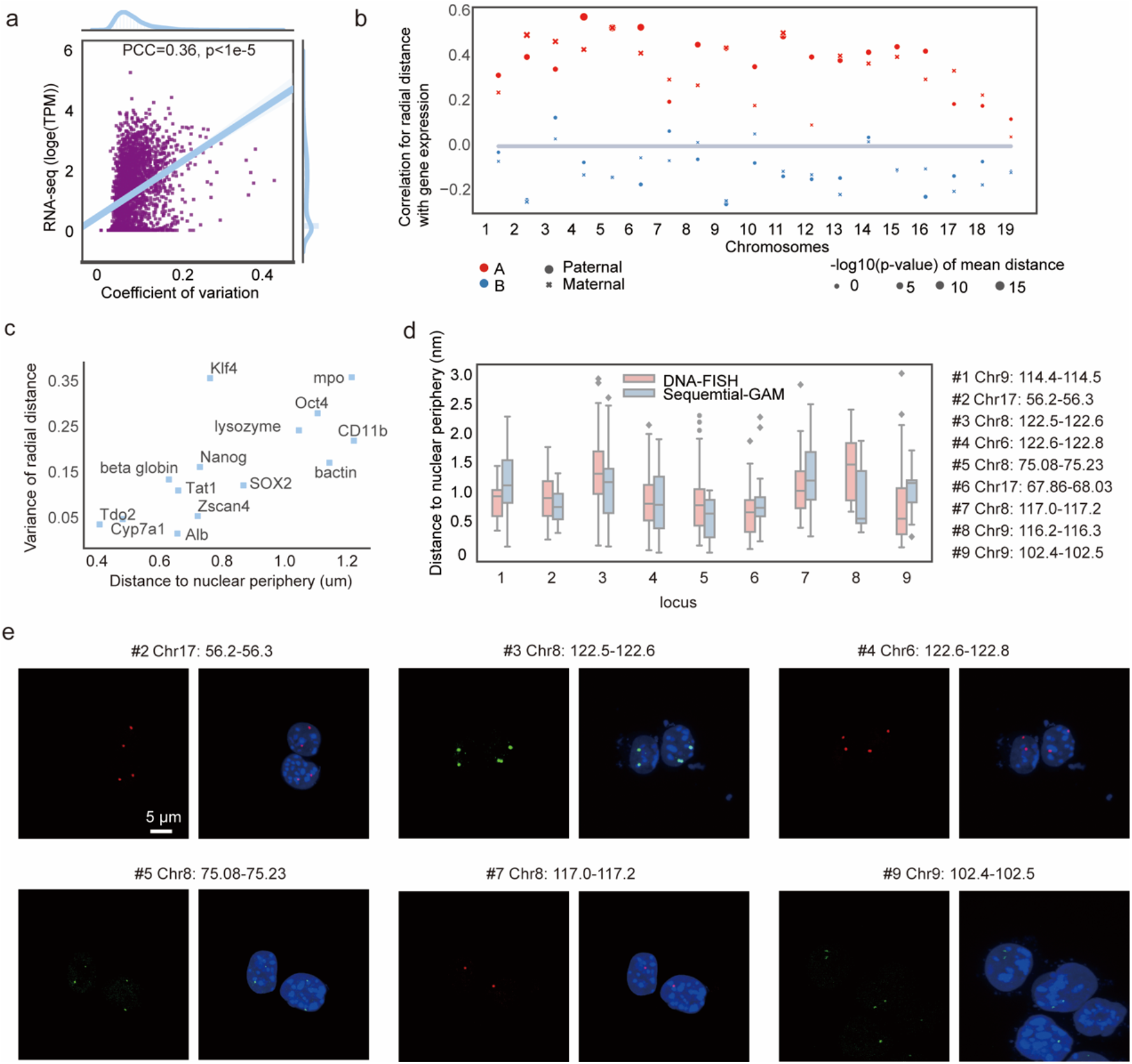
The radial distribution of genes correlates with their local status. **(a)** Correlation between gene expression and the variance of radial distance (Pearson correlation coefficient = 0.36, p-value <1e-5), with the linear correlation shown in the blue line and the 95% confidence coefficient shown in the blue shadow. Each dot here represents one 1Mb locus, with 2416 loci in total. **(b)** Correlation between gene expression the variance of radial distance of loci belonging to compartment A (red) or B (blue) in each chromosome on paternal (dot) and maternal (cross) genome. **(c)** Examples of the radial position and the variance of active and inactive genes in mouse ES cells. Red spots mark active genes, and blue spots mark repressed genes in mouse ES cells. **(d)** Comparison of loci distance to nuclear periphery in DNA-FISH and Sequential-GAM. The p-values of Student’s t test between DNA-FISH and Sequential-GAM are all larger than 0.01, showing no significant difference. **(e)** The representative DNA-FISH results. The green signals (Alexa-488) and red signals (Alexa-647) represent DNA loci. Blue signals in the middle represent nuclei after DAPI staining.

We further selected nine genes to validate the above results, including highly expressed genes and inactive genes in mouse ESC. Gene loci of OCT4, CD11b, SOX2, NANOG, Klf4, β-actin in mouse ESC exhibit smaller distances to the nuclear center than the inactive genes such as TDO2, TAT1, CYP7α, ALB, β-globin (Fig. 6c), which were similar to the position of these gene in previous study^20^. The radial position of nine genomic loci was further validated by DNA-FISH (Fig. 6d, see Methods). The distances of distinct loci to nuclear periphery in Sequential-GAM were consistent with those in DNA-FISH (Fig. 6e, Supplementary Table S5).

Overall, these results indicate that gene expression levels decrease and structural stability improves progressively from the nuclear center to the periphery, whereas the association between expression and radial distance, and the association between expression and structural stability, decreases gradually.

## Discussion

In this work, we developed Sequential-GAM to investigate 3D genome structure in single diploid cells. Through a series of slices in one cell, geometric structure in a single cell could be obtained by Sequential-GAM. We found that chromosome location was correlated with averaged gene density. Compartment A and active histone marks preferred locating at the center of the nucleus, while compartment B and inactive histone marks preferred positioning at the nucleus periphery. The radial location of subcompartments A1, A2, B1, B2, and B3 was gradually positioned from the nuclear center to the periphery, but with dynamics in the order to the center of nuclear. The radial distributions of histone modifications and subcompartments were in accordance with the gradient model; that was, active and inactive chromatin formed a continuous gradient along the nuclear radius, as supported by previous studies^14,38,39^. The q-stable TADs were identified, and the relationship between chromatin spatial structure stability and gene expression was investigated. The results demonstrated that the stability of q-stable TADs was correlated with the enrichment of LADs, the deletion of particular genes, and the number of subcompartment species. Finally, we found a substantial relationship between the genes’ radial positions and expression levels, and variations with their expression levels. In summary, by constructing the allelic 3D genome structure of single cells, Sequential-GAM contributes to the understanding of the genome’s dynamics and stability.

Comparing to other methods, Sequential-GAM and GAM are more powerful to detect long-range interactions than Hi-C and SPRITE^40^. Moreover, Sequential-GAM supplies the geometric insight into the genome.

The q-stable TADs in single mES cells were defined with a miniature spatial variance among TADs. Similarly, TAD cliques in Hi-C have been reported, mainly in heterochromatin around the stable nucleus^34^. The TAD cliques, based on topological inter-TAD interactions (Supplementary Fig. 9g), reflected topologically condensed structures, while q-stable TADs revealed stable structure between TADs in 3D nuclei. In q-stable TADs, the link between stability and expression was affected differently by the varied roles of the genes. One restriction might be that the paternal and maternal genomic readings were identified using SNPs of CAST/129S4. 129S4 has significantly fewer SNPs than CAST (about 1/6 times), resulting in an imbalance of parental data. For example, the q-stable TADs in 129S4 were much smaller in CAST due to losing part of 129S4 data (Supplementary Fig. 9).

There are some limits for Sequential-GAM. For example, the limited genome capture rate in Sequential-GAM could come from the experimental procedure. In Sequential-GAM, we identified consequent slice from single-cell under 150X-objective microscope, which may cause relatively low efficiency of genome amplification and low quality because of long exposure in the environment^41^. This can be improved by the recently developed enhancement of micro-DNA sequencing technology^42^. Furthermore, the cell number in this study is limited. In Sequential-GAM, the process of tracing continuous thin sections, laser dissection and whole genome amplification was done from cell to cell, slice by slice. Normally, it costs about two days and 1200 dollars per nucleus if 20 continuous slices were captured in one nucleus, so there is no second cell type shown but other cell lines can be explored in the future by Sequential-GAM.

Moreover, part of the public data (e.g., the RNA-seq data) utilized in this study were for bulk sample, while the results of geometric feature were based on the ten mES cells we captured, which may lead to an incomplete description of the overall distribution of chromatin. To compensate for this, we obtained statistically significant results by increasing the amount of data. For examples, in the exploration of the relationship between gene expression and radial stability, the results were made by 2416 loci (Fig. 6a) with significant p-value. As the technology develops and more samples become available, researchers will be able to explore more solid geometric information.

Besides, since Sequential-GAM employed a sphere for nucleus approximation, for other shapes of cells, the relevant shape (e.g., ellipsoidal) must be at first mapped into the sphere, then computed under constraints, and then mapped back to the original shape to create the structure accurately. Additionally, although we did not calculate the geometric features between chromosomes in the main text, Sequential-GAM could be extended to calculate the relative positions between chromosomes (Supplementary Methods).

Last but not the least, the previous imaging-based sequencing methods revealed the genome-wide chromatin structure, RNA expression levels, and chromatin topologies^41,43^. It will be very promising to combine Sequential-GAM with genome-wide immunolabeling in tissues, to obtain higher dimensional single-cell genome structure *in situ*.

## Online Methods

### Cell culture

The mouse embryonic stem cell F123 cell, described previously^44^, was a generous gift from Prof. Wei Xie’s laboratory. F123 cell line was F1 hybrid from *Mus musculus castaneus* × S129/SvJae. The cells were grown in DMEM medium containing 15% FBS (Gibco), 1×penicillin/streptomycin (Gibco), 1× non-essential amino acids (Gibco), 1× GlutaMax (Invitrogen), 1000 U/mL mLIF (Millipore, ESG1107), 0.4mM β-mercaptoethanol. F123 cells were initially cultured on feeder cells (inactivated MEF (mouse embryonic fibroblast) cells. Feeder MEF cells were derived from the E13.5 ICR mouse. MEF medium is DMEM within 10% FBS, 1×penicillin/streptomycin, and 1× GlutaMax. MEF cells were inactivated by X-ray (3.5 Gy/mL). Before F123 cells were used, cells were cultured on 0.1% gelatin-coated feeder-free plates for two passages or sorted by FACS by Hoechst 33342 staining.

### Cell staining and G1 phase sorting

F123 cells were harvested in single-cell suspension and fixed by 4% and 8% depolymerized (EM-grade) paraformaldehyde for 10 min and 2 h, respectively. Then cells were stained with 1μg/ml Hoechst 33342 or not (negative control), incubated at 37°C for 20 min, keeping dark. Then cells were washed once with PBS, and resuspended in PBS. 10^6^ G1 phase cells were sorted at excitation 353nm (BD Aria SORP).

### Sequential cut by ultrathin cryosection and laser dissection

The protocol for cells preparation and ultrathin cryosection, referred to the Tokuyasu method, was employed with some modification^45^. Briefly, sorted G1 phase F123 cells were harvested and fixed by 4% and 8% depolymerized (EM-grade) paraformaldehyde for 10 min and 2 h, respectively. Then cells were resuspended and pelleted in 1% gelatin to remove extra water. Then, cells were resuspended in 12% gelatin and solidified adequately at 4°C. After 30min, the gelatin within cells was sliced into approximately 0.5 mm^3^ cubes and selected according to cell density under stereoscope. The cubes were embedded in saturated 2.1 M sucrose overnight before freezing in liquid nitrogen. The frozen cell cube was cryosectioned into 200nm-thickness slices and placed on PET membrane slide (Leica, 11505151) through loop with sucrose. Particularly, consequent thin sections were collected from the top to the bottom of the frozen cell cube. The slices on membrane slide were rinsed with sterilized nuclease-free water gently and air-dry. Under 150 × microscopy, the continuous nuclear slices from the same cell were selected and laser dissected with laser (LMD7000), collected into the lid of PCR tubes containing cell lysis buffer.

### Whole Genome amplification (WGA)

Collected single nuclear slices in Cell 3-10 were amplified using MALBAC single cell genome amplification kit (Yikon, KT110700150), following manufacture instructions. Generally, laser dissected nuclear slice was lysed, reverse cross-linked by proteinase K and fragmented at 50°C for 50 min following 80°C for 10 min. Next, DNA was pre-amplified when supplied with 30 μL pre-Amp buffer and 1 μL pre-Amp Enzyme under the gradually increasing temperature 20°C for 40sec, 30°C for 40sec, 40°C for 30sec, 50°C for 30sec, 60°C for 30sec, 70°C for 4min, 95°C for 20sec, and finally 58°C for 10sec. Collected single nuclear slices in Cell 1-2 were amplified using single-cell whole genome amplification kit (WGA4, 254-457-8), following manufacture instructions.

Generally, laser dissected nuclear slice was lysed, reverse cross-linked by proteinase K and fragmented at 50°C for 4h. Next, DNA was fragmented and amplified. Then, DNA was amplified after adding 30 μL Amplification buffer and 0.8 μL Amp enzyme. Amplified DNA was purified using 0.75 × volume of VATHS DNA beads (Vazyme, N411-03) following manufacture instructions. Then, DNA was amplified after adding 30 μL Amplification buffer and 0.8 μL Amp enzyme. In addition, 1 ng purified whole genomic DNA or several live cells acts as positive controls. 1μL molecular water or 10 μm diameter empty PET membrane were negative controls. The controls were proceeded as same as the above samples. Amplified DNA can be stored at -20 °C until proceeding further library preparation.

### Library construction and sequencing

The sequencing libraries for each slice of nuclear were constructed using TruePrep DNA Library Prep Kit from Illumina. 5ng of amplified DNA was used for library construction, following the manufacturer’s instructions (Vazyme, TD502). The concentrations of libraries were estimated using Qubit 2.0 fluorometer (Thermo Fisher Scientific). Library size distribution and pooling were performed using Agilent 2100. Each nuclear profile was sequenced in pair-end 150 bp on Illumina Hiseq 2000. We obtained 2 batches of data with 10 cells for the experiments in total. Cell1-4 is for the 1^st^ batches, and Cell 5-10 is for the 2^nd^ batches.

### DNA-FISH and Imaging data processing

Briefly, pre-plated F123 cells on 0.1% gelatin were fixed by 4% paraformaldehyde for 10 min following permeabilization. Next, cells were treated with 0.1 M HCl for 30 min and incubated in 50% formamide/2×SSC for 10 min. A prepared fluorescent probe was added into cells and incubated at 75°C for 5min and followed 37°C for12 hours in a moist chamber (keep dark). Then cells were washed and stained with DAPI. Air-dried coverslips were mounted on glass slides. Images were acquired on a confocal microscope and distances between the center of FISH signals and nuclear periphery was measured in Imaris 9.6 and the detailed data was seen in Supplementary Table S5.

### Noise filtering

First, we calculated the distribution of the number of reads for different windows of the positive chain at each resolution (10kb, 100kb, 1Mb) by *bedtools coverage*. Then, we calculated the 0.1 quantile of the overall distribution of reads, and also the 0.1 quantile of the distribution of reads for each nuclear profile. As a result, we set 1 as the threshold of the number of reads in the filtered windows. That is, we filtered the windows with the number of reads less than or equal to 1 in each nuclear profile. Finally, we kept the corresponding negative chain reads according to the remaining positive chain reads, and the two were combined as the result after noise filtering.

### Determining the resolution

For each DNA fragment, its genomic start *G*_*start*_was mapped to a different resolution (e.g., *R*_1000_if the resolution is 1000 bp), in order to determine its coordinate (e.g., *G*_1000_ if the resolution is 1000 bp) after the change in resolution. That is:

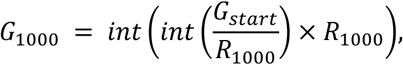

in which the *int* means rounding down. For example, location 101,126,000 bp would be mapped to 101,000,000 for resolution of 1Mb.

After changing the resolution, the genome coverage (*C*_*g*_) of each cell at several resolutions (1kb, 10kb, 100kb, 1000kb) was computed.

In addition, we estimated the theoretical capture rate (*C*) for each cell based on the number of nuclear slices captured (*N*_*np*_), the radius of the nucleus (*R*), and the slice thickness (*L*_*cut*_). That is, regarding the capture rate to be proportional to the volume captured, and the volume captured was roughly estimated using the ratio of cut thickness to diameter:

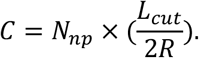

As depicted in Supplemental Figure S3g, we compared the theoretical DNA capture rate to the genome coverage (*C*_*g*_) for each resolution. The result indicated that at a resolution of 10 kb, the two parties demonstrated were in agreement. At resolutions greater than 10 kb, the capture rate for each fragment was overestimated, whereas resolutions less than 10 kb did not account for experimental and sequencing losses. Consequently, 10kb was selected as the final resolution.

### Determining the parents of the reads

Since the CAST/129S4 cell line was chosen for Sequential-GAM, the parentage of the fragment could be determined by SNP.

Using the sam2seg function of the Dip-C tool^46^, we determined whether a parental or maternal SNP was present for each DNA fragment based on the SNP data (data source is in http://renlab.sdsc.edu/huh025/snp/), and subsequently labeled the fragments accordingly. In this step, some DNA fragments that lacked SNP information were not labeled as parental or maternal.

Next, we use the K nearest neighbors (KNN) algorithm (*neighbors*.*KNeighborsClassifier* in the sklearn package for python) to classify the DNA fragments (*R*_*unknown*_) that are unsure of their parentage based on their genomic coordinates 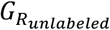 *and* the z-axis information 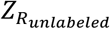 of the nucleus fragment where the DNA fragment is located. KNN is utilized because DNA fragments belonging to the same chromatin are more likely to be located close to one another in space.

Specifically, we initially train the KNN model using DNA fragments labeled with parental information 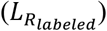 with genomic coordinates as 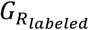 and z-axis information as 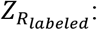

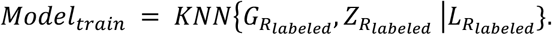

The trained model is then used to classify unlabeled DNA fragments based on information from the five samples surrounding each sample:

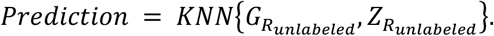

Consequently, for each genomic segment, its corresponding parental marker is obtained.

To determine the z-axis positions of the reads from the parent and the fragments from the mother at 10kb resolution, we mapped the DNA reads for each parent to the locus at 10kb resolution. As an estimate of the z-axis coordinates for this 10kb resolution window, we calculate the average of the z-axis coordinates of the DNA fragments in windows containing multiple DNA reads.

### Segmentation of fragments located in LAD

If the processed 10 kb windows intersect with the cLAD (overlap > 1bp), then the fragment is considered to be at the edge of the nucleus. Since the reference coordinate of the cLAD data in the publicly available data is mm9, we used the *liftover* tool in the UCSC Genome Browser to map the coordinates to mm10 before the calculation.

### Radius estimation of nuclear profile

As a result of Fluorescence-activated cell sorting, the cells in Sequential-GAM are of uniform size. Therefore, we divide the sorted cells into two groups and use one group to estimate the nucleus-to-cytoplasm ratio of the second group.

One group of the cells was stained separately for nucleus and cytoplasm, and the ratio of nucleus to cytoplasm in around 100 cells in suspension was calculated based on the staining results. As a result, the average nucleus-to-cytoplasm ratio was 0.7, and the nucleus radius was 4.3975um.

For the other group of cells, we performed the Sequential-GAM slicing experiment and independently measured the radius of each slice (which is radial of cell but not radial of the nuclear). We obtained an estimate of the nuclear radius for each slice *r*_*1*_, *r*_*2*_, *⋯, r*_*n*_ by multiplying by 0.7. By recording the order of cutting, we also obtain the distance *d*_*1*_, *d*_*2*_, *…, d*_*n*_ between each nuclear profile (200 nm between adjacent slices), which is used to calculate the z axis coordinates of each nuclear profile.

### Calculation of z-axis coordinates

Since the nucleus was not sliced from top to bottom, the z-axis coordinates *z*_*1*_, *z*_*2*_, …, *z*_*n*_ of each slice were undetermined. As there was an error in the radius of the nucleus for each slice, using the radius of a single slice to estimate the nucleus’ z-axis would result in a larger error. To reduce the overall error for a given cell, we aligned the radius of all its slices with the cell’s location along the z-axis (Supplementary Fig. 6h). The specific details are listed below.

Given the radii *r*_*1*_, *r*_*2*_, …, *r*_*n*_ and interval distances *d*_*1*_, *d*_*2*_…, *d*_*n*_, the following conditions must be fit for *z*_*1*_, *z*_*2*_, …, *z*_*n*_, which aimed to find the best z-aixs locations to reduce the error:

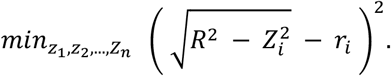

Consequently, the z-axis position of each slice was computed under the condition of minimizing this equation.

### Sequential-GAM computational method

Assuming that the nucleus is spherical, a column coordinate system was established with the center of the nucleus sphere as the coordinate origin as in the figure below:

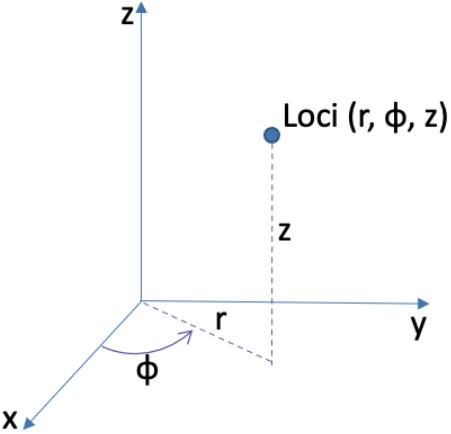

Then the coordinate of the loci in the column coordinate system was *loci*_*i*_ *= (r*_*i*_, *ϕ*_*i*_, *z*_*i*_).

Correspondingly in the right-angle coordinate system, the coordinate was:

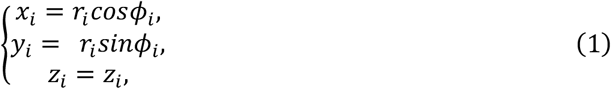

From the location of given nucleus profiles (NPs), we determined the z-axis information where each locus was located, then the z-axis coordinates were known for all loci.

We defined the power-law equation between genome distance (*x*) with spatial distance (*y*) as:

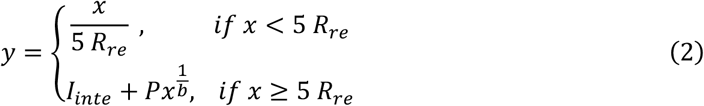

where *R*_*re*_ is the resolution, *I*_*inte*_ is the intercept, *P* is the coefficient for power-law, b is the value of the power (sample from the distribution calculated from data of DNA seqFISH+^16^). For distinct cells and chromosomes, we collected samples of distinct parameters.

All loci *{S*_*all*_} were divided into two sets *{S*_*l*_*} and {S*_*n*_}, in which *{S*_*l*_} represents the loci located at cLAD, and *{S*_*n*_} represents the loci not located at cLAD. The cLAD data for different cells were sampled at a rate of 90% from the bulk data.

For any two *loci*_*i*_, *loci*_*j*_ with supposed coordinate (*x*_*i*_, *y*_*i*_, *z*_*i*_) and (*x*_*j*_, *y*_*j*_, *z*_*j*_), respectively. By the genome distance *G*_*ij*_, the theoretical spatial distance 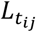 could be obtained according to equation (2). Besides, their spatial distance 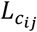 was estimated as follow:

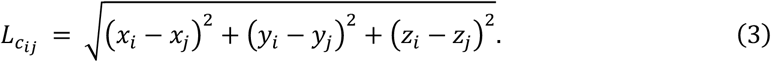

Then Sequential-GAM calculated the error *RES*_*)l*_ between the estimated spatial distance and the theoretical spatial distance for all loci:

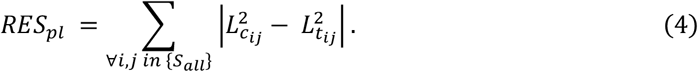

Using the genome distance *G*_*ij*_ *as t*he normalization factor, the constrain *C*_*pl*_ of power-law for the loci is:

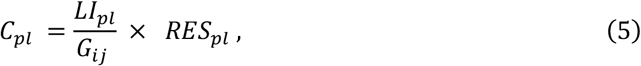

where *LI*_*pl*_ is the constrain parameter for power-law for calculation (1e-7 in our experiments. This value affects the speed of convergence, but not the final convergence result).

Next, the distance between the loci and the cLADs was used to constrain the position of the loci. For a given *loci*_*i*_, Sequential-GAM found the nearest cLAD loci in the genome distance and returns the genome distance as *G*_*i-LAD*_. *G*_*i-LAD*_ was put into equation (2) to yield *d*_*i-LAD*_, representing the theoretical distance of the loci to the nucleus’s edge.

Then, the estimated distance of loci from the edge of the nucleus is the radius of the nucleus *R minu*s its distance to the center of the sphere *r*_*i-center*_:

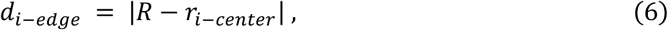

where *r*_*i-center*_ is calculated according to the coordinates of this locus in the column coordinate system *(r*_*i*_, *ϕ*_*i*_, *z*_*i*_), which is 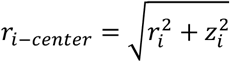.

Then the error between the estimated distance from loci to cLAD and the theoretical distance to the edge of the nucleus is:

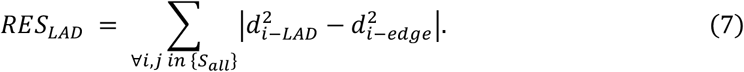

Using the genome distance *G*_*i-LAD*_ as the normalization factor, the constrain *C*_*LAD*_ of cLAD for the loci is:

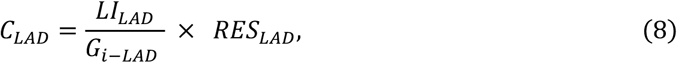

where *LI*_*LAD*_ is the constrain parameter for LAD (1e+9 in our experiments). The *LI*_*LAD*_ do not affect the final result of convergence.

Finally, the loss function is the sum of the two constrain (5) and (8):

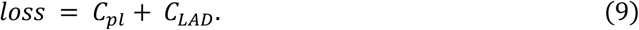

The final estimate of each loci coordinate is obtained by optimizing the loss function (9). Due to the different degree of capture of each chromosome, we calculated each chromosome individually and did not perform an overall optimization for all chromosomes of the same cell.

### Sphericity, volumes, and surface area to volume ratio

In this manuscript, the principal axis length of chromatin refers to the length of the long axis of the ellipsoid obtained by fitting these points with an ellipsoid, using the points of the captured chromatin as samples.

The volume of chromatin refers to the volume occupied by the convex polyhedron formed in the frame of the captured points of chromatin. The surface area of chromatin refers to the surface area of the corresponding convex polyhedron.

Sphericity refers to the degree to which the chromatin is close to a sphere. In this paper, the sphericity of chromatin is calculated from the volume and surface area of chromosomes as follows.

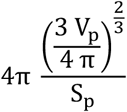

where V_p_ is the chromosome volume, and S_p_ is the chromosome surface area. When the chromosome is spherical, the sphericity is equal to 1.

### Calculation of distance matrix

After calculating the spatial location of each 10kb locus, we calculated the two-by-two Euclidean spatial distances between them to obtain the 10kb resolution distance matrix. As the distance matrix for other resolutions, such as the 1Mb resolution, the average distance between the 10kb loci contained within each 1Mb bin was subsequently calculated. The parental genomes showed different coverage because of the slices’ position and SNPs. The position of captured slices in the same cell may not cover the same proportion in two parental alleles. In F123 cells, the number of SNPs in paternal genome is larger than that in maternal genome. Therefore, we calculated the 3D structure and distance matrix separately for the parents. When generating the contact matrices (distance matrix), for the accuracy of the results, we did not use interpolation to fill in the missing loci in this version. For single cells, we used blanks for the uncaptured parts. For the average results of multiple cells, we only calculated the average of captured loci.

### Calculation of TADs and q-stable TADs

TAD boundaries were calculated by the *hicFindTAD* tool from HiCExplorer^47,48^ in mES cells at 40kb resolution, and in total of 6215 TADs. The location of each TAD was estimated based on the mean location of all captured loci within it. Then the distance matrix M_d_ was calculated by the Euclidean distance between each two TADs. In addition, the variance matrix M_v_ of the TAD distance matrix between the 10 cells was derived from the variation of this distance matrix in 10 cells, i.e. the distance variation between the *i*^*th*^ *and j*^*th*^ *TAD* is *m*_*vij*_. We referred the mean of variation matrix M_v_ as mean _*mv*_, and the standard deviation of variation matrix M_v_ as std_*mv*_. After filtering to obtain TAD pairs with high confidence (details are in the Supplementary) in M_v_, each pair of TADs satisfied *m*_*vij*_ < mean_*mv*_ *™ std*_*mv*_ was the relatively stable TAD pairs, as a set *S*_*stable*_

The TAD cluster analysis was then performed. Then, we constructed a graph consisting of TADs as vertices and variance as edges (keeping only TAD pairs in *S*_*stable*_). Using the Louvain algorithm^49^, we computed the partition of the graph nodes, which maximizes the modularity. The TADs are relatively stable within each modularity (the result of clustering), whereas the distance between each modularity is unstable. This modularity is a candidate for quasi-stable TADs. A cluster here is a candidate quasi-stable TADs set.

To further identify the stable portion of the candidates, the coefficient of difference (details are in the Supplementary) of the q-stable TADs was calculated, and the portion with a coefficient of difference smaller than the median of all the candidates is referred to as the quasi-stable TADs set (termed q-stable TADs).

### Obtaining of the most stable part and relatively unstable part of q-stable TADs

To determine the extreme portion of q-stable TADs, the 0.3 quantile and 0.7 quantile for all q-stable TADs were first calculated. Then, the q-stable TADs with coefficient of difference less than 0.3 quantile were set as the most stable part, and those with coefficient of difference greater than 0.7 quantile were set as the relatively unstable part.

### Dividing of the q-stable TADs with the content of specific genes or regulators

First, we calculated the 0.3 and 0.7 quantile of peaks of specific genes or regulators (house-keeping genes, OSN genes, super-enhancer and LADs) by the ChIP-seq data (data sources are in the Supplementary Table S3) of binding sites. Thereafter, we denote the q-stable TADs with peaks number less than 0.3 quantile as the depletion group, the q-stable TADs with peaks number not less than 0.7 quantile as enrichment group, others as random group.

### Stability of the radial positioning of genes

For each gene, with a 1Mb locus in its neighborhood in each cell, we calculated the mean location for the 1kb loci we captured as the location of the 1Mb locus in Euclidean space. Then, we calculated the radial distance between the location and the center of the nucleus. Regarding the stability, the standard deviation of the radial distance from the nucleus center in ten cells per 1Mb locus was computed. The greater the variance in 1Mb locus positioning, the lower the structural stability.

## Acknowledgements

We thank Prof. Wei Xie from Tsinghua University, and Prof. Minping Qian from Peking University for the helpful discussion and suggestions. We thank Dr. Ying Li (Cryo-EM Facility of Tsinghua University, Branch of National Protein Science Facility) and the imaging facility of Tsinghua University for the technical support. We thank the team member Mr. Yan Yan from Prof. Michael Zhang, student Ms. Leyuan Meng from JiLin University, China, and Mr. Haochen Su from Cambridge University, UK for part of FISH experiment preparation. This work was supported in part by National Key projects for international cooperation on science, technology and innovation (2021YFE0201100), the National Natural Science Foundation of China (81890991) and Beijing Municipal Natural Science Foundation (Z200021) to J.G., National Nature Science Foundation of China (62050152) to M.L.S., and CAS Interdisciplinary Innovation Team (JCTD-2020-04) to J.G.. M.Q.Z acknowledges the support by the Cecil H. and Ida Green Distinguished Chair.

## Conflicts of interest

The authors declared no competing interests.

## Author contribution

Conceptualization: Yongge Li, Kaili Wang, Michael Zhang, Juntao Gao; data generation: Kaili Wang, Yongge Li; data curation: Yongge Li, Kaili Wang; formal analysis: Yongge Li, Kaili Wang; funding acquisition: Michael Zhang, Juntao Gao; investigation: Kaili Wang, Yongge Li, Michael Zhang, Minglei Shi, Juntao Gao; methodology: Yongge Li, Kaili Wang, Michael Zhang; project administration: Michael Zhang, Juntao Gao, Minglei Shi; resources: Michael Lab; software: Yongge Li, Kaili Wang; supervision: Michael Zhang, Juntao Gao, Minglei Shi; validation: Yongge Li, Kaili Wang, Juntao Gao; visualization: Yongge Li, Kaili Wang; writing: Kaili Wang, Yongge Li, Michael Zhang, Minglei Shi, Juntao Gao

## Data availability

The data is in NCBI with number GSE193989.

## Code availability

The code of Sequential-GAM is in https://github.com/yonggeli66/Sequential-GAM.

## Notes

### Competing Interest Statement

The authors have declared no competing interest.

